# RAPPPID: Towards Generalisable Protein Interaction Prediction with AWD-LSTM Twin Networks

**DOI:** 10.1101/2021.08.13.456309

**Authors:** Joseph Szymborski, Amin Emad

**Affiliations:** Department of Electrical and Computer Engineering, McGill University, Montréal, QC, Canada; Mila, Quebec AI Institute, Montréal, QC, Canada; The Rosalind and Morris Goodman Cancer Institute, Montréal, QC, Canada

## Abstract

**Motivation:** Computational methods for the prediction of protein-protein interactions, while important tools for researchers, are plagued by challenges in generalising to unseen proteins. Datasets used for modelling protein-protein predictions are particularly predisposed to information leakage and sampling biases.

**Results:** In this study, we introduce RAPPPID, a method for the Regularised Automatic Prediction of Protein-Protein Interactions using Deep Learning. RAPPPID is a twin AWD-LSTM network which employs multiple regularisation methods during training time to learn generalised weights. Testing on stringent interaction datasets composed of proteins not seen during training, RAPPPID outperforms state-of-the-art methods. Further experiments show that RAPPPID’s performance holds regardless of the particular proteins in the testing set and its performance is higher for biologically supported edges. This study serves to demonstrate that appropriate regularisation is an important component of overcoming the challenges of creating models for protein-protein interaction prediction that generalise to unseen proteins. Additionally, as part of this study, we provide datasets corresponding to several data splits of various strictness, in order to facilitate assessment of PPI reconstruction methods by others in the future. Availability and Implementation: Code and datasets are freely available at https://github.com/jszym/rapppid.

**Supplementary Information:** Online-only supplementary data is available at the journal’s website.

## INTRODUCTION

Interactions of proteins with other proteins and their surroundings are fundamental to the internal machinery of a cell. These interactions are of particular interest, as it is essential for a bevy of diverse cellular functions: from organising cell structure to generating metabolic energy (Huttlin *et al*., 2017). These interactions are typically validated with a high degree of confidence by the many biological assays commonly employed today, each with their own specific advantages and challenges (Snider *et al*., 2015). Assays for validating protein interactions range from the venerable yeast two hybrid (Y2H) (Vidal and Fields, 2014) which researchers have relied on for the past decades, to more recent Biotin-related techniques such as BioID-MS (Roux *et al*., 2012). A characteristic of all these assays, however, is that they are costly in terms of time, labour, and materials. A further complication of protein-protein interaction studies are the multiple sources of biases that plague small and large datasets alike. The choice of proteins to include in a study poses a particular threat of so-called “bait bias” in smaller datasets, while large datasets suffer from biases in the discovered protein interactions (“prey bias”) as well as the exacerbation of laboratory biases when included in aggregated datasets (Gillis *et al*., 2014). Entire classes of proteins, such as membrane proteins, which are sometimes difficult to experimentally validate are often under-represented in these interaction studies as well.

Computational approaches to predict protein-protein interactions (PPIs) are therefore useful to help towards reducing the number of costly experiments researchers are required to perform. Researchers have deployed many diverse approaches to solve the task of protein sequence-based interaction prediction. Most sequence-based methods rely on the understanding that co-evolution and co-expression of proteins are both tied to protein interaction and sequence similarity (Cong *et al*., 2019; Jansen, 2003). Some methods rely on substitution matrices for sequence alignment such as BLOSUM or PAM in combination with machine learning methods to predict interactions (Henikoff and Henikoff, 1992; Ding *et al*., 2016). Other methods utilise Support Vector Machines (SVMs) with kernels specifically designed for use with protein sequences (Ben-Hur and Noble, 2005). Other statistical methods including naÏve bayes (NB) and *k*-nearest neighbors (kNN) have been used to predict protein interactions from protein sequences (Browne *et al*., 2007). Some of the most successful PPI prediction methods belong to the family of methods that rely on novel substring search algorithms (Li and Ilie, 2017; Dick *et al*., 2020), operating similarly to sequence search tools like BLAST (Altschul *et al*., 1990). Deep learning models have also been designed for predicting protein interactions (Chen *et al*., 2019). These deep approaches commonly either learn wide networks or share weights in a twin design; the latter being shown to be both more efficient and effective (Richoux *et al*., 2019).

Such methods, however, face many challenges due to the nature of the data on which they train. Arguably the most pervasive of which is the ability of models to generalise and predict the interactions of proteins previously unseen by the prediction method. To ensure such generalisability, careful cross-validation techniques must be used to avoid data leakage. While the necessity of appropriate cross-validation techniques is not unique to this area of research, the application of these networks (e.g., PPIs, transcriptional regulatory networks) to obtain biological insights makes it particularly important to address this challenge in the task of biological network reconstruction (Park and Marcotte, 2012; Tabe-Bordbar *et al*., 2018).

For PPI reconstruction, developing generalizable models prove particularly difficult. The nature of PPI networks makes it easy to create datasets with testing/training splits which leak information, resulting in inflated performance metrics that cannot properly assess the generalisability of these methods. In particular, simply splitting interaction datasets into training and testing sets using random selection of edges results in the construction of testing datasets that are almost entirely comprised of interactions between proteins found in the training set. Indeed, Park and Marcotte in 2012 found that all the PPI prediction models they surveyed were tested on such naïvely constructed datasets (Park and Marcotte, 2012). The surveyed models, which had optimised their performance on these naÏve datasets, that suffered from a large degree of information leakage, were also found to incur precipitous falls in their prediction metrics when tested on datasets where no proteins in the testing set occurred in the training dataset.

When faced with obstacles in the construction of generalisable models, strategic and targeted applications of regularisation at training time can significantly improve results. This is particularly relevant in the context of deep learning methods which pay for their expressiveness by learning an outsized number of parameters. Among the most common regularisation techniques used in the deep learning context is “dropout” (Srivastava *et al*., 2014). Applying dropout to a layer consists of randomly zeroing the activations of the previous network with some probability *p*. However, choice of regularisation techniques must be selected with care and in accordance with the architecture. For example, applying dropout directly to the hidden state of Recurrent Neural Networks (RNNs) (Lipton *et al*., 2015), an architecture which lends itself naturally to sequential inputs such as amino acid chains, impairs its ability to retain its memory of previous inputs (Zaremba *et al*., 2015).

Applying regularisation to RNNs requires additional considerations. Recent work by Merity *et al*. has demonstrated that randomly zeroing the weights of RNNs (“dropconnect”) (Wan *et al*., 2013) rather than their hidden state activation (“dropout”) effectively reduces testing error (Merity *et al*., 2017). Merity *et al*. describe applying dropout to the embedding layer as well as using an averaged optimiser (NT-ASGD) as part of a series of regularisation techniques dubbed Averaged Weight-Dropped Long Short-Term memory (AWD-LSTM). The regularisation techniques used by AWD-LSTM models are specifically selected for their suitability in the context of training RNNs.

To meet the generalisation challenges posed by PPI prediction tasks, we developed a method called the Regularised Automatic Prediction of Protein-Protein Interactions using Deep Learning, or RAPPPID. RAPPPID addresses the challenges in creating generalised models for PPI prediction by adopting (with modification) the AWD-LSTM, a regularised recurrent neural network training routine (Merity *et al*., 2017). In this study, we showed that RAPPPID outperforms state-of-the-art PPI prediction methods on strict validation datasets constructed in accordance with guidelines set out by Park & Marcotte (Park and Marcotte, 2012). Additionally, we performed various analyses and applied RAPPID to different use-cases to show-case the effect of different components of its architecture and training on its performance and its applicability to different real-world scenarios.

It is worth mentioning that in addition to RAPPPID’s code, we have made various pre-processed datasets freely available in the hope that they facilitate the evaluation of protein interaction prediction methods on datasets that both mitigate information leakage and are appropriately large for deep learning applications.

## METHODS

### PPI prediction using AWD-LSTM Twin Networks

RAPPPID is first trained by considering pairs of amino acid sequences of proteins along with a label indicating whether they do or do not interact. The amino acid sequences are first tokenised using the Sentencepiece algorithm (Kudo and Richardson, 2018), which allows for better recognition of common groupings of amino acid residues that make up the secondary structure and motifs of proteins. Fixed-length latent vector representations of the token sequences of both proteins are then computed by twin neural networks (Bromley *et al*., 1993), forming the encoder of RAPPPID’s pipeline. These twin networks have shared architectures and weights and are trained jointly. An overview of the pipeline is provided in Figure 1.

**Figure 1.**
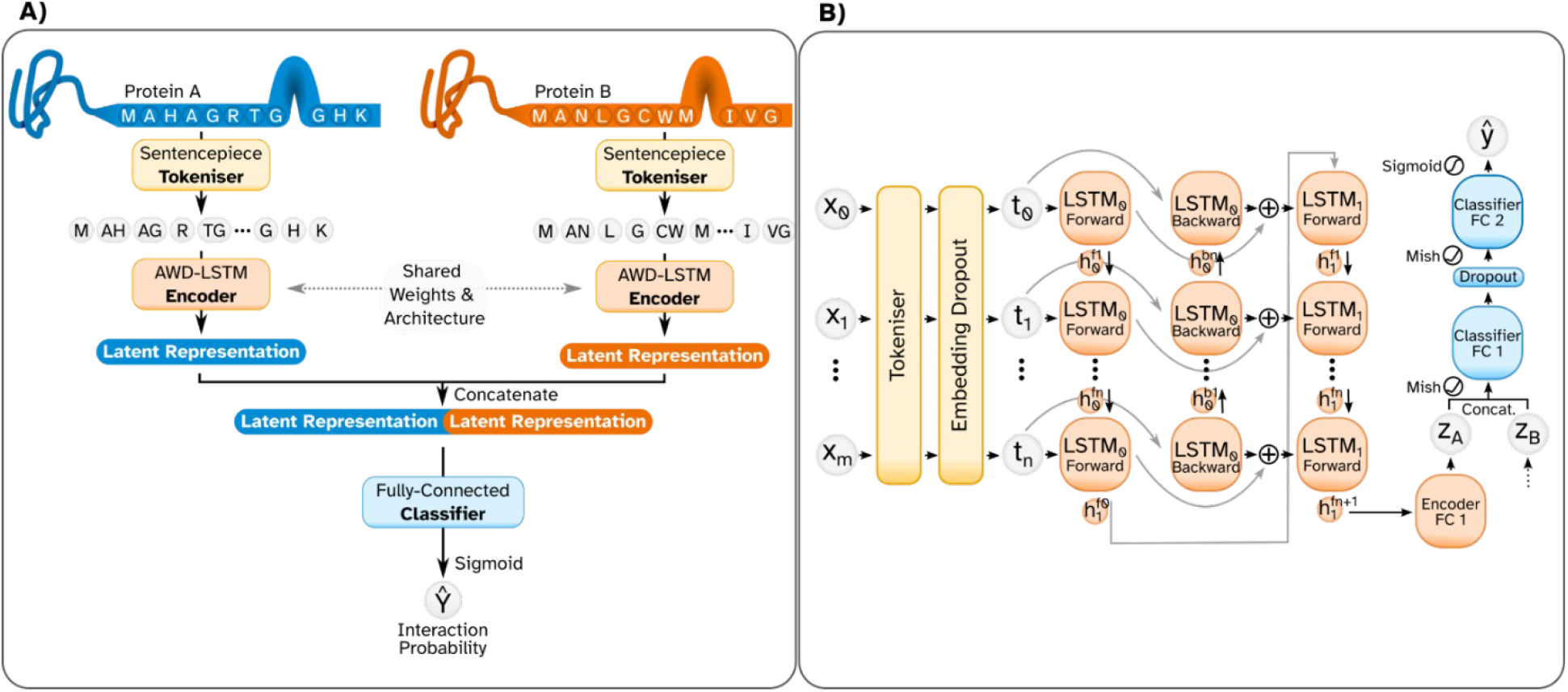
Overview of the RAPPPID pipeline and architecture. (A) The pipeline of the RAPPPID begins with two protein sequences. Each sequence is first tokenised by the Sentencepiece tokeniser (yellow). Each sequence of tokens is then inputted separately into an AWD-LSTM encoder layer which results in a latent representation for each protein. The latent representations of each protein are inputted into a fully-connected classifier layer. The classifier layer outputs the predicted probability of the two proteins interacting. (B) Taking a closer look at the architecture of the RAPPPID, individual residues *x*_0_ to *x*_*m*_ for a protein comprised of *m* residues are tokenized into *n* tokens *t* _0_, *t* _1_,*…, t* _*n*_. The embedding dropout layer randomly assigns random tokens from the total vocabulary to zero. The encoder layer is comprised of a multi-layer bidirectional LSTM whose last hidden state is fed to a fully connected layer before outputting a latent representation *z* _*A*_. *z* _*A*_ is then concatenated with the latent representation of a second protein (*z*_*b*_) before being inputted into the two-layer fully-connected classifier. The output of the classifier is activated by the sigmoid function to produce a probability of interaction.

Each twin network consists of a two-layer bidirectional AWD-LSTM network (Merity *et al*., 2017), which takes a tokenised amino acid sequence as its input and generates a fixed-length latent vector representation as its output. AWD-LSTMs are architecturally identical to the LSTMs (Hochreiter and Schmidhuber, 1997), however at inference time, several regularisation techniques are employed while training to promote learning generalised weights. Among these regularisation techniques are an averaged optimizer and dropout applied to embeddings and LSTM weights (Athiwaratkun *et al*., 2019; Wan *et al*., 2013). Using an AWD-LSTM encoder enables RAPPPID to leverage the strong inductive biases of LSTMs, while ensuring that the learned weights are generalised.

The final hidden state of the AWD-LSTM is passed to a single fully-connected layer whose output, once activated by the Mish function (Misra, 2020), is the latent representation of the amino acid sequences. The latent representations of both proteins are then concatenated and provided as inputs to the classifier network which generates an interaction probability for each protein pair. The classifier network is a two-layer fully-connected network that outputs a single logit whose sigmoid activation serves as the probability of the two proteins interacting. The activated logit is then used to calculate the mean binary cross-entropy loss. Relegating the pairwise comparison of proteins to the shallower classifier network allows RAPPPID to infer protein interactions in an efficient manner. Figure 1 provides an overview of RAPPPID’s pipeline.

### Sequence Segmentation and Tokenisation

As mentioned earlier, RAPPPID utilises the Sentencepiece algorithm (Kudo and Richardson, 2018) to tokenize amino acid sequences. While words form the basis of many natural languages and may break up sentences and phrases into discrete units, no such higher-order segmentation is as immediately apparent in amino acids. Motifs and protein domains possess many analogous qualities to words in natural languages; they appear repeatedly in amino acid sequences and their combination and relative position in these sequences play important roles in the protein structure and function (Anfinsen, 1973).

Much like words, however, motifs and protein domains present an “out-of-vocabulary” problem, where unseen examples are difficult to handle. Attempts to solve this problem in natural language processing tasks has resulted in “subword” segmentation algorithms, particularly in difficult-to-segment languages such as Japanese which do not separate words by spaces (Schuster and Nakajima, 2012). Here, we employ the Sentencepiece algorithm (Kudo and Richardson, 2018) to sample tokens from “subword” vocabularies generated by the Unigram algorithm (Kudo, 2018). The unigram and sentencepiece algorithms construct vocabularies of arbitrary size by modelling the probabilities of subwords and provides a principled manner for sampling from this distribution to reconstruct sequences. The multi-residue tokens that comprise the vocabulary subdivide low-entropy areas and reduce the overall length of the sequences encoded.

### Generalising Protein Sequence Encoding with AWD-LSTM

The task of protein-protein interaction prediction on unseen proteins is a difficult problem prone to overfitting, as demonstrated by the poor testing performance of various methods on unseen proteins (Park and Marcotte, 2012). For this reason, a training and optimisation methodology that allows efficient regularisation is desirable. AWD-LSTM was recently devised to enable efficient training of generalisable recurrent neural networks (RNNs) (Merity *et al*., 2017). This approach deploys several regularisation techniques during training to achieve this goal. RAPPPID adopts, with modification, the training methodology of AWD-LSTM.

RAPPPID utilises Embedding Dropout, DropConnect (Wan *et al*., 2013), and Weight Decay (Loshchilov and Hutter, 2019) on the LSTM weights, as described by AWD-LSTM, in the encoder during training. Both AWD-LSTM and RAPPPID optimise over average weights, however the optimisers are quite different. AWD-LSTM makes use of the non-monotonically triggered averaged stochastic gradient descent (NT-SGD) optimiser which switches between stochastic gradient descent (SGD) and the averaged variant (ASGD). RAPPPID uses the recent Stochastic Weight Averaging (SWA) strategy in combination with the Ranger21 optimiser (Athiwaratkun *et al*., 2019; Wright and Demeure, 2021).

SWA has been shown to promote generalisable models in part by overcoming the challenges of finding best solutions within flat loss basins (Izmailov *et al*., 2019). Ranger21 is an optimiser that applies the “Lookahead mechanism” (Zhang *et al*., 2019) to the AdamW optimiser (Loshchilov and Hutter, 2019) and includes several optimisation techniques (Wright and Demeure, 2021). These techniques, which include gradient centralisation and adaptive gradient clipping, enable us to further improve our ability to learn generalised weights and smooth training trajectories (Brock *et al*., 2021; Yong *et al*., 2020). Finally, since RAPPPID does not rely on the timestep outputs of the LSTM network, the Temporal Regularisation (TAR) described by AWD-LSTM is not applicable.

### Details of RAPPPID’s architecture and hyperparameter tuning

The dimensionality of the AWD-LSTM hidden state is made equal to the dimensionality of the embeddings (*i*.*e*., 64). The output of the fully-connected layer is of equal dimensionality to that of the AWD-LSTM hidden state and is activated by the Mish activation function (Misra, 2020). The output of the first fully-connected layer is half the size of the embedding dimension, activated by the Mish function, and regularised by a dropout layer (Srivastava *et al*., 2014). RAPPPID trains on a vocabulary of 250 tokens that is generated by the Sentencepiece algorithm. New vocabularies are generated for each dataset before training.

The number of LSTM layers, L2 coefficient, and various dropout rates are defined as hyper-parameters that require tuning between different datasets. Hyper-parameters were selected by cross-validation. The range of considered hyperparameters and their selected values are provided in the Supplementary Tables S1-S2 (in Supplementary File 1). The chosen hyper-parameter ranges are a compromise between common, reasonable values and maintaining a manageable hyper-parameter space which can be explored practically.

### Protein Interaction Datasets

We obtained protein-protein interactions (PPIs) and protein sequences from version 11 of the Search Tool for the Retrieval of Interacting Genes/Proteins (STRING) database (Szklarczyk *et al*., 2019), from the official STRING website. Edges were downloaded from https://stringdb-static.org/download/protein.links.detailed.v11.0.txt.gz and sequences were downloaded from https://stringdb-static.org/download/protein.sequences.v11.0.fa.gz. In this dataset, the association between any two proteins is assigned a confidence score depending on the source of the information (called a “channel”). In our analyses, only associations with a combined STRING-score above 95% (equivalent to a score above 950, obtained from combining different channels) were considered as positive edges. We also retained the channel-specific scores for further analysis.

To obtain an estimate of the false-positive rate of the confidence score-filtered STRING dataset, we leveraged the curated and experimentally validated non-interacting protein pairs from the Negatome dataset (Blohm *et al*., 2014). By comparing the set of proteins that are in both STRING and Negatome, and evaluating the number of negative edges in Negatome that were considered a positive edge in this intersection, we estimated the false-positive rate of our STRING dataset to be 4.01%. This false-positive rate is within the expected 5% upper-bound given by our 95% confidence threshold.

PPI graphs are understood to be scale-free in the general case. This property of PPI graphs can make them challenging datasets upon which to train a generalised model, as some proteins can be over-represented. To characterise the extent to which proteins are represented in the dataset, we calculated the distribution of the relative degree of proteins in the network (Supplementary Figure S1A-C). The vast majority of proteins (upwards of 85%) have edges with fewer than 1% of proteins within their dataset split (*i*.*e*., train/validation/test). The protein with the highest degree is CDC5L which has a relative degree of just over 10%, while the second highest has a relative degree of 6.2% (Supplementary Figure S1D-F).

### Negative Examples

The preparation of datasets of pairs of proteins which are known not to interact with one another is a fraught process. Various methods are typically deployed to create such datasets, which are integral for machine learning methods as they typically require negative examples. Ideally, only negative examples are entirely composed of non-interacting protein pairs which are experimentally verified and manually curated, such as the Negatome database (Blohm *et al*., 2014).

Unfortunately, non-interacting pairs are difficult to experimentally validate. As a result, there are far fewer *H. sapiens* negative protein pairs in such datasets such as Negatome than positive protein pairs in databases such as STRING. Precisely, there are 1,191 negative *H. sapiens* pairs in Negatome, and 263,130 positive pairs above a 95% confidence threshold in STRING (only pairs comprised of proteins present in UniprotKB were included in this analysis). The resulting class imbalance is detrimental to learning performant, generalisable models.

Synthetic negative pairs are often used to compensate for the small number of experimentally verified non-interacting pairs is construction of synthetic negative pairs. A common method for constructing negative pairs is to select pairs of proteins from distinct sub-cellular compartments. This method has unfortunately been shown to result in biased samples according to multiple measures (Ben-Hur and Noble, 2006). Selecting pairs at random from the space of pairs not known to interact has proven to evade such biases, but runs the risk of capturing yet unknown false-negatives. Unless otherwise noted, RAPPPID utilises synthetic random pairs of proteins which are not known to interact.

### Training, Validation and Testing Set Construction

As identified by Park and Marcotte (Park and Marcotte, 2012), methods which consider the interaction of proteins in a pairwise fashion must (and have historically failed to) take additional care to avoid information leakage when constructing training and testing datasets. Following their suggestion, here we use three different classes of testing and training sets to evaluate the performance of RAPPPID and other algorithms (Figure 2).

**Figure 2.**
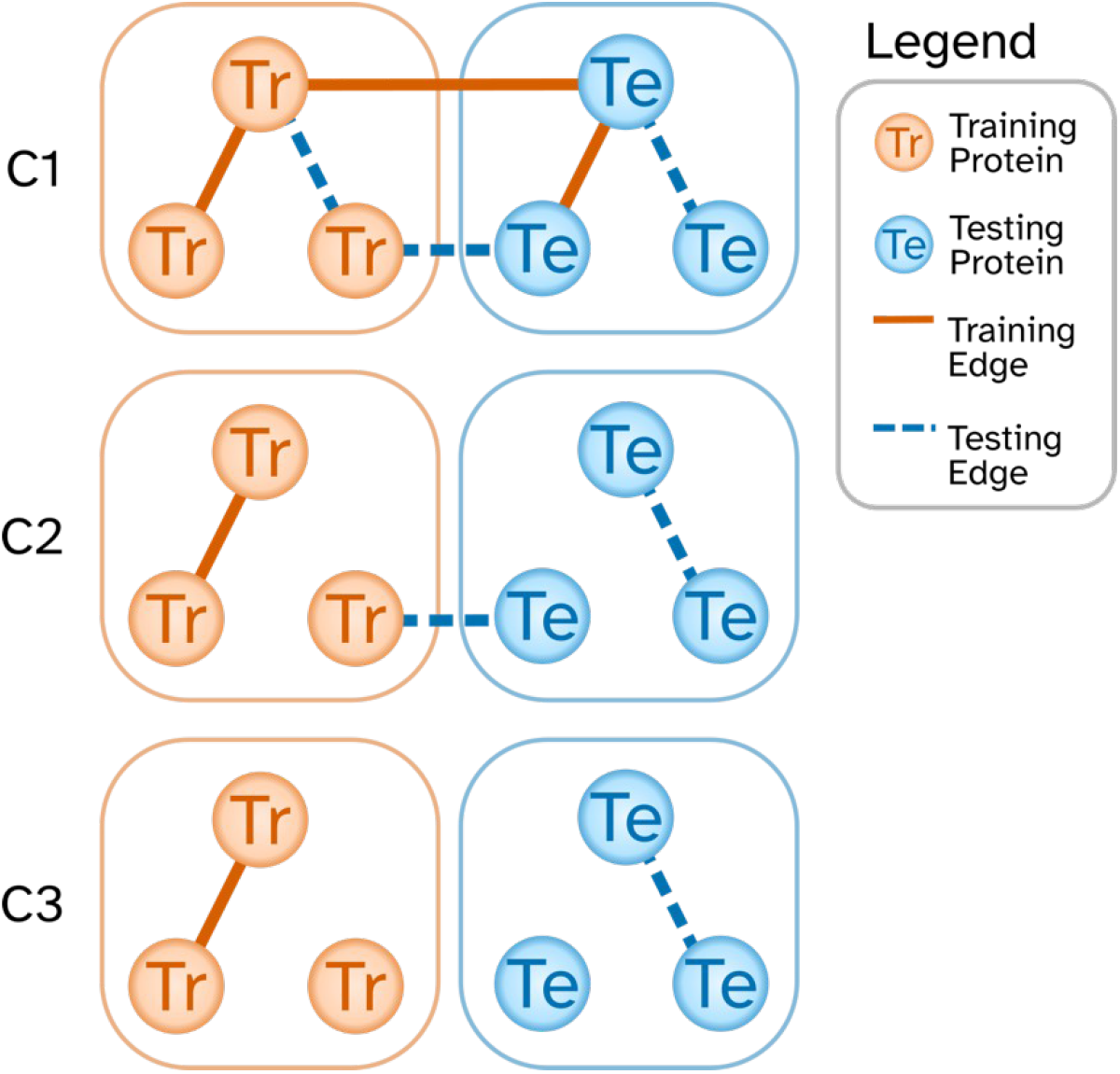
Illustration of differences in edges between C1, C2, and C3 datasets. Differences between C1, C2, and C3 datasets are most visible by first dividing the population of all proteins in the dataset into training (orange, left) and testing (blue, right). In the case of the strict C3 dataset (bottom row), edges known at training time (orange, solid) only occur between training proteins. Similarly, C3 datasets are evaluated on testing edges (blue, dotted) that only occur between testing proteins. The C2 dataset has all the edges present in the C3 dataset, but also includes testing edges between testing proteins and training proteins. Finally, the pervasive C1 datasets allows all possible training and testing edges that are not identical.

1. “C1” refers to the evaluation scheme in which edges (pairs of proteins) are randomly selected to form the training or testing sets. Since the selection criterion is based on edges, both proteins in the pair may be present in both the testing and training sets (due to the presence of other edges adjacent to each protein).
2. “C2” refers to the evaluation scheme in which proteins are randomly selected to form the training or testing sets. In this scheme, only one protein in a pair may be present in both testing and training sets (but never both). This evaluation scheme mimics the scenario in which a model trained on the interactome is used to predict the interaction of known proteins with a newly discovered protein (that was not used to train the model).
3. “C3” refers to the evaluation scheme in which proteins are randomly selected to form the training or testing sets. However, unlike C2, proteins which appear in the training set never appear in the testing set. This is the most strict evaluation scheme.

While most methods have typically reported on models validated with datasets in the C1 class, they often perform much worse on similar datasets in the more conservative C2 and C3 class. This is likely due to the information leakage between training and testing sets present in the C1 class and, to a lesser extent, in the C2 class. In our evaluations, we report the performance of different methods using all three evaluation schemes above, but we are most interested in the results of C3 due to a lack of information leakage.

In the C-type datasets we’ve created, there are approximately 9.3 thousand proteins. The number of edges in the datasets range from just over 263 thousand edges in the C1 dataset to just under 174 thousand edges in the C3 dataset. Full dataset statistics are presented in Supplementary Table S3. These C-type datasets are made freely available to the public, with instructions on how to download them at https://github.com/jszym/rapppid/tree/main/data. We’ve made these datasets available in the hope that it facilitates the evaluation of protein interaction prediction methods on datasets that both mitigate information leakage and are appropriately large for deep learning applications. The adoption of C-type datasets for training and evaluation of PPI prediction methods is important for accurate and representative benchmarking.

### Implementation

RAPPPID was implemented in the Python computer language using the PyTorch and PyTorch Lightning deep learning framework (Paszke *et al*., 2019; Falcon *et al*., 2020). Embedding Dropout and DropConnect implementation were obtained from the AWD-LSTM code base (Merity *et al*., 2017). The source code for RAPPPID can be found by visiting https://github.com/jszym/rapppid. RAPPPID was trained at a rate of approximately 2.7 training steps per second at a batch size of 80 protein pairs on an NVIDIA RTX 2080 GPU and 32 CPU cores clocked at 2.2 GHz. The C3 model takes approximately 2.4 hours to train, while C1 and C2 take approximately 7.8 hours.

### Protein Similarity Experiments

In the analysis of the sequence similarity between testing and training proteins, “Percent Identity” was measured between proteins using NCBI’s PSI-BLAST tool running locally as part of version 2.12.0 of NCBI’s BLAST+ software suite (Altschul *et al*., 1997). In addition to the Percent Identity, for two proteins to be considered similar, an E-value cut-off of at most 5 and an alignment length of more than 30% of the query sequence were considered necessary. The 64-bit Linux binaries of the BLAST+ suite were obtained from the link ftp.ncbi.nlm.nih.gov/blast/executables/blast+/2.12.0/.

## Results

### Performance evaluation of RAPPPID and other algorithms

To establish the ability of RAPPPID to correctly predict protein-protein interactions within the current landscape of PPI prediction methods, we compared it against three recent methods (Figure 3). The first of these is the Scoring Protein INTeractions (SPRINT) method, which belongs to the family of methods that predict interactions according to measures of sequence similarity (Li and Ilie, 2017). SPRINT was shown to outperform support-vector machine (SVM), random-forest (RF), and sequence similarity-based methods across C1, C2, and C3-like datasets.

**Figure 3.**
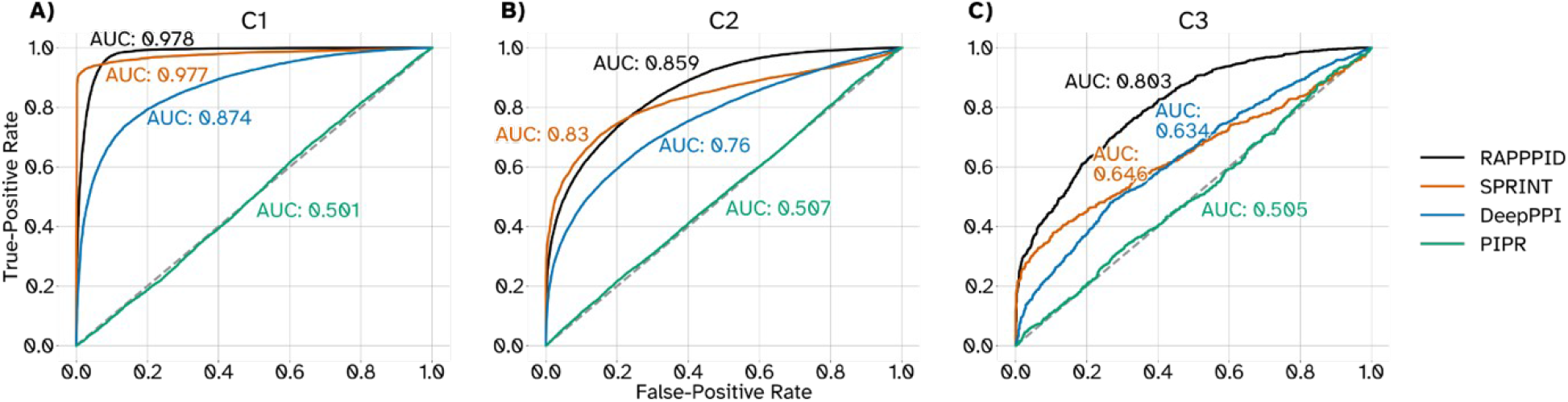
Receiver-Operator curves across methods and datasets. The receiver-operator curves (ROCs) for all four methods tested across C1 (A), C2 (B), and C3 (C) datasets.

The two other methods, PIPR and DeepPPI, are deep learning methods that similar to RAPPPID utilize twin networks (Chen *et al*., 2019; Richoux *et al*., 2019). PIPR uses a residual recurrent convolutional neural network (RCNN) for its encoder with the goal of more effectively summarising both local and global features. We compared RAPPPID against the best performing iteration of DeepPPI, whose encoder comprises of a convolutional neural network feature extractor followed by an LSTM network. All three methods required retraining, as the available weights of these models belong to diverse datasets, all of which are either not C2 or C3 type, or (in the case of SPRINT) are insufficiently large for training a generalisable deep learning model.

Across C1, C2, and C3 testing datasets, RAPPPID achieved higher area under the receiver-operator curve (AUROC) than all other methods tested (Table 1). The margin between RAPPPID and the second highest performing method (SPRINT in all cases) was highest when performed on the stricter C3 dataset, resulting in approximately a 24.3% improvement. The improvement obtained by RAPPPID compared to SPRINT was lower on the C2 dataset (approximately 3.4%), and finally nearly equivalent on the least strict C1 dataset.

**Table 1:** Comparison of PPI prediction performance on C1, C2, and C3 datasets. The testing AUROC and AUPR of four different PPI prediction methods is reported across the three different dataset types described by Park & Marcotte (Park and Marcotte, 2012).

| Dataset | Method | Testing AUROC | Testing AUPR |
| --- | --- | --- | --- |
| C1 | RAPPID | **0.978** | 0.974 |
|  | SPRINT | 0.977 | **0.983** |
|  | DeepPPI | 0.874 | 0.881 |
|  | PIPR | 0.501 | 0.405 |
| C2 | RAPPID | **0.859** | **0.868** |
|  | SPRINT | 0.830 | **0.868** |
|  | DeepPPI | 0.760 | 0.787 |
|  | PIPR | 0.507 | 0.508 |
| C3 | RAPPID | **0.803** | **0.810** |
|  | SPRINT | 0.646 | 0.716 |
|  | DeepPPI | 0.574 | 0.590 |
|  | PIPR | 0.505 | 0.509 |

**Table 3:** Results from an ablation study conducted on RAPPPID. Each model is trained/tested twice on three randomly generated C3 datasets. The performance metrics correspond to held-out test sets.

|  | RAPPPID<br>(original) | RAPPPID-<br>SWA | RAPPPID<br>+Adam | RAPPPID-<br>AWD | RAPPPID-SentencePiece | RAPPPID<br>+TransfLG | RAPPPID<br>+TransfSM |
| --- | --- | --- | --- | --- | --- | --- | --- |
| **Test AUROC** | 0.792<br>($\pm 0.007$) | 0.782<br>($\pm 0.007$) | 0.791<br>($\pm 0.025$) | 0.762<br>($\pm 0.020$) | 0.749<br>($\pm 0.009$) | 0.670<br>($\pm 0.030$) | 0.747<br>($\pm 0.026$) |
| **AUROC Diff** | N/A | -1.20% | -0.100% | -3.70% | -5.37% | -15.3% | -5.68% |
| **Test APR** | 0.794<br>($\pm 0.009$) | 0.783<br>($\pm 0.007$) | 0.792<br>($\pm 0.032$) | 0.757<br>($\pm 0.022$) | 0.748<br>($\pm 0.011$) | 0.686<br>($\pm 0.040$) | 0.758<br>($\pm 0.025$) |
| **APR Diff** | N/A | -1.37% | -0.273% | -4.62% | -5.85% | -13.6% | -4.61% |

With regards to the area under the precision-recall curve (AUPR), this trend across dataset types persisted. RAPPPID’s AUPR was higher than all other methods for the C3 dataset, with a margin to the second highest method of 0.094 (equivalent to an approximately 14.6% improvement). RAPPPID’s AUPR score was matched by SPRINT in experiments conducted on the C2 dataset, but outperformed DeepPPI and PIPR. SPRINT achieved the highest AUPR of all the methods in the C1 dataset, outperforming RAPPPID with a margin of 0.009 (equivalent to an approximately 0.9% improvement).

While we were able to replicate results of the PIPR model on the *S. cerevisiae* dataset published as part of the original PIPR publication (Chen *et al*., 2019), PIPR suffered from convergence issues during training on our *H. sapiens* STRING datasets. We suspect this is due to a variety of factors, with the large differences in dataset characteristics being the most likely cause. The number of proteins and interactions in the *S. cerevisiae* dataset is far smaller than our STRING datasets. Furthermore, the *S. cerevisiae* dataset selected pairs of proteins which occupy different subcellular compartments in order to construct negative samples; an approach found to lead to biased estimations of prediction accuracy (Ben-Hur and Noble, 2006).

Taken together, these results suggest that RAPPPID outperforms alternative methods in the majority of evaluations on C1, C2, and C3 schemes and is particularly effective in the stricter and more difficult C3 evaluation.

### Channel-specific performance of RAPPPID

The STRING database, integrates and annotates protein association data from a wide range of sources. The “database”, “text-mining”, “experiments”, and “coexpression” channels make-up the majority of the edges in our datasets (e.g., 98.4% of all the edges in the C3 dataset).

The “database” channel is comprised of several curated databases of interactions such as KEGG and Reactome (Kanehisa, 2000; Jassal *et al*., 2019). Edges in the “text-mining” channel are the result of a statistical analysis of proteins whose names and/or identifiers co-occur in publications. The “experiments” channel is populated by interactions evidenced by high-throughput experiments curated by members of the International Molecular Exchange (IMEx) consortium (Orchard *et al*., 2012). This includes datasets such as IntAct, DIP, the BioGRID, and many others (Orchard *et al*., 2014; Salwinski *et al*., 2004; Oughtred *et al*., 2020). Finally, the “coexpression” channel arises from proteomic and transcriptomic assays which quantify gene-by-gene correlations. STRING additionally assigns calibrated confidence scores to each of the edges which summarise the evidence supporting an edge. Scores are assigned to each edge by channel, and finally harmonised into a final “combined confidence score” which represents the evidence across all channels present in STRING.”

To better characterise the results of our protein-protein prediction tests, we sought to identify the source of the testing edges RAPPPID correctly and incorrectly identified. Figure 4A and Supplementary Figure S2 show that RAPPPID can accurately predict the testing set edges that have a high confidence score in biologically supported channels of co-expression, experiments, and database. However, the accuracy for the edges that have a high confidence score in the text-mining channel is inferior to the other channels. Since edges that are only supported by text-mining (but not by the other channels) are arguably the ones most prone to error, we expected RAPPPID to have an inferior performance on such edges (since the edges themselves may not be reliable). To test whether the inferior performance of RAPPPID in the text-mining channel in Figure 4A are indeed due to such edges, for a fixed threshold k (50 ≤ k ≤ 95), we divided the testing edges with a text-mining confidence score at least equal to k into two groups: a group with “experiments” confidence score at least equal to 80% and a group with “experiments” confidence score smaller than 80% (Figure 4B). Evaluating the testing edges in C1, C2, and C3 showed that for this channel, the accuracy on the former group is higher than the latter group (sometimes as large as ∼22% higher, Supplementary Figure S3). Repeating the analysis for co-expression and database channels also confirmed this trend (Figure 4B, Supplementary Figure S3). Taken together, these results suggest that the inferior performance of RAPPPID on the text-mining channel in Figure 4A is indeed due to the edges that are supported only by text-mining and not by other biologically identified channels.

**Figure 4.**
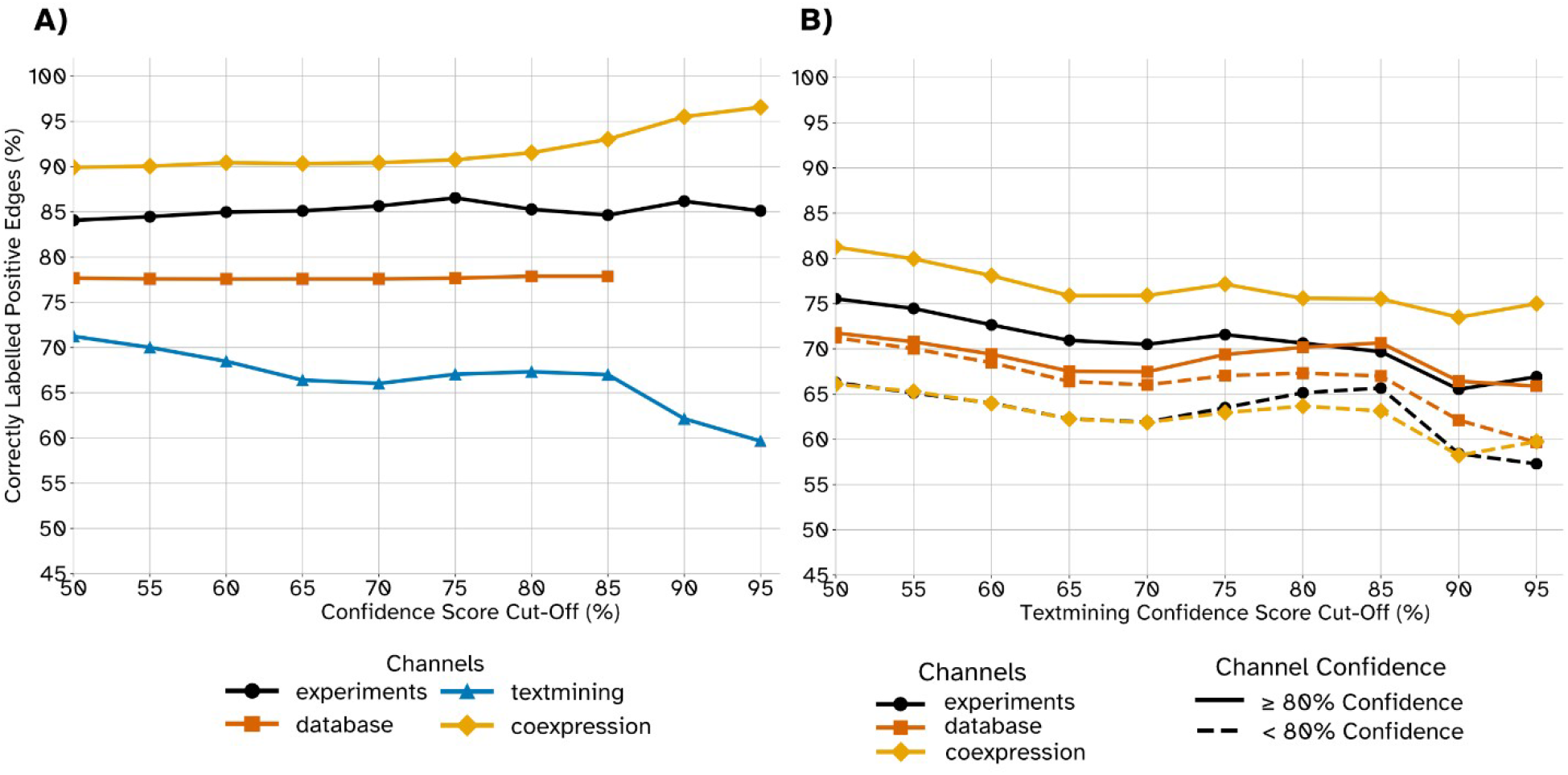
Accuracy of positive edges across edge confidence stratified by STRING channels. (A) The percentage of correctly labelled positive edges are plotted for each major STRING channel. The x-axis denotes the channel edge confidence cut-off score for each curve’s respective channel. (B) Here, we see a similar chart but rather than using each channel’s respective score as a confidence cut-off, edges are excluded according to their text-mining confidence (x-axis). The solid curves include edges which have a channel confidence *≥* 80 % for the channel indicated by the curve’s colour. Dashed curves conversely include edges whose channel confidence is ¿ 80 % for the channel indicated by the curve’s colour. In both (A) and (B) data shown reflects the C2 model/dataset.

### Role of Protein Similarity on RAPPPID’s Performance

The procedures for C2 and C3 datasets were devised to reduce the information leakage by avoiding testing on edges which contain proteins which are known to an algorithm during its training. This safeguard against information leakage, however, does not account for proteins which are known by different identifiers but share near identical protein sequences. Further complicating the matter, sequence similarity is a valid PPI prediction feature and is often used by methods as a proxy measure of co-evolution and conserved functional domains (Cong *et al*., 2019; Jansen, 2003). Indeed, this strategy is leveraged by the SPRINT algorithm against which RAPPPID was compared.

In spite of the challenges above, we sought to determine whether the superior performance of RAPPPID (particularly in the strict datasets of C2 and C3) is due to sequence similarity between testing and training proteins or not. For this purpose, we used PSI-BLAST algorithm (Altschul *et al*., 1997) to evaluate sequence similarities between each pair of testing/training proteins. Figure 5 shows the accuracy of RAPPPID on the C2 dataset when different degrees of restriction on sequence similarity are imposed. More specifically, a threshold *t* (x-axis in Figure 5) determined the maximum allowable Percent Identity score between a testing protein and any of the training proteins that were candidates to be similar to it (see Methods for details). Any testing protein that did not satisfy this condition for the threshold *t* was excluded from the calculation of accuracy. As one moves towards larger values of *t*, the sequence similarity constraint loosens and *t*=100% is equivalent to the complete C2 dataset. Our analysis on C2 (Figure 5) and C3 (Supplementary Figure S4) revealed that RAPPPID’s accuracy is largely independent of the sequence similarities between testing and training proteins and the performance of RAPPPID does not deteriorate when removing testing proteins that have a highly similar training protein.

**Figure 5.**
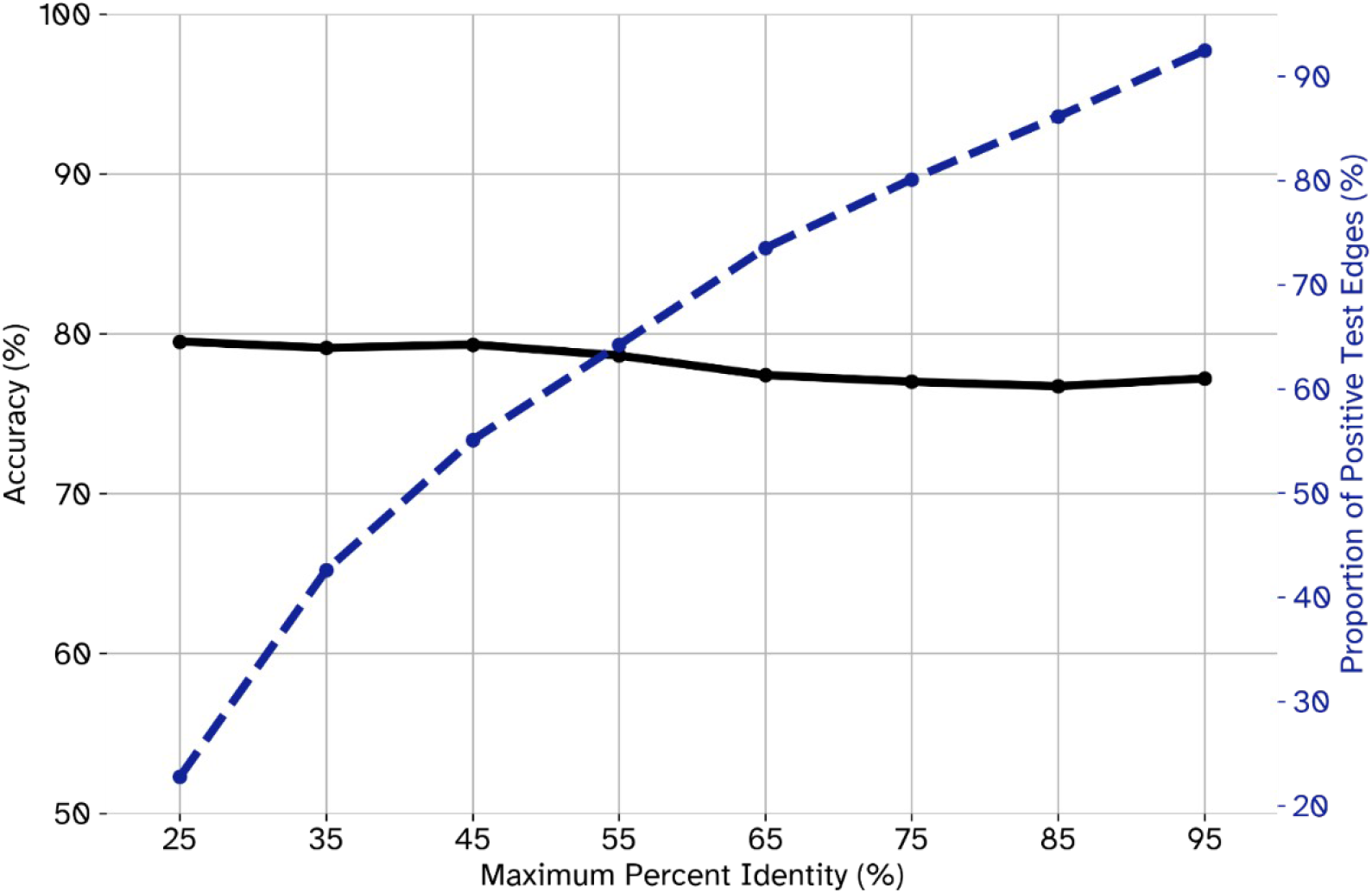
Accuracy of positive edges as a function of similarity between testing and training proteins in C2. The similarity between testing and training proteins was measured using their percent identity as computed by NCBI’s PSI-BLAST software. The highest percent identity between any training protein and a testing protein in a testing edge was considered to be that testing edge’s “maximum percent identity”. The percentage of accurately labelled positive edges (black curve, left y-axis) is reported for edges with maximum percent identities lower than the threshold reported on the x-axis. The proportion of testing edges for each threshold values is reported by the dashed blue curve and the right y-axis.

### Effect of different components on RAPPPID’s performance

To understand RAPPPID’s performance when trained and tested on negative examples which are experimentally validated and manually curated, we created a new C3 dataset where the negative examples are provided by the Negatome database of non-interacting pairs. To combat the large class imbalance caused by the relatively few pairs of human proteins in Negatome (see Methods for details), additional random pair negatives were supplemented until both positive and negative classes had an equal number of samples. After 13 epochs, RAPPPID achieved a testing AUROC of 0.802 and a testing AUPR of 0.799 on C3 dataset, which is comparable to the performance of RAPPPID using randomly selected negative edges.

Since RAPPPID utilizes random initialisation, mini-batch sampling, and token sampling, there is some stochasticity present in its performance. First, we sought to test the effect of these stochastic components and to ensure that the specific choice of training and test sets were not responsible for the superior performance of RAPPPID. For this purpose, we ran RAPPPID twice on three additional C3 datasets whose training, validation, and testing proteins were chosen at random (a total of six times). In each of these runs, different seeds were used to assess the effect of stochastic components of RAPPPID (Table 2). Overall, the average testing AUROC across models trained on these additional three datasets was 0.792 (±0.007). Results from these repeatability experiments illustrate that RAPPPID’s strong performance on C3 datasets is not tied to a specific set of testing or validation proteins, nor certain weight initialisation states.

Next, we conducted an ablation study (Table 2) using the same three C3 datasets and two runs of each variant per dataset (a total of six models per variant). Each variant substitutes one component of RAPPPID’s regularisation, architecture or optimisation, allowing us to quantify the difference in performance for which these components are responsible. In this table, “RAPPPID-SWA” variant removes SWA, “RAPPPID+Adam” replaces Ranger21 with Adam optimizer (learning rate selected by hyperparameter tuning), “RAPPPID-AWD” removes the AWD regularisation, and “RAPPPID-SentencePiece” substitutes the SentencePiece tokens with amino-acid-residue-level tokens. As can be seen in this table, the choice of optimizer has negligible effect on the performance. However, SWA, AWD, and SentencePiece tokens all contribute to the superior performance of RAPPPID, with the tokeniser having the largest contribution. One advantage that SentencePiece affords RAPPPID is the regularising effect of random sampling tokens. Additionally, the SentencePiece tokeniser reduces the total length of sequences inputted into the tokeniser. The protein sequences on which RAPPPID is trained, while limited to 1,500 residues, are sufficiently large enough for gradients to vanish quiet substantially; a phenomenon long known to plague RNNs when trained on long sequences (Hochreiter, 1998).

In acknowledgment of the recent prevalence and performance of attention-based methods in deep learning (Vaswani *et al*., 2017), the sixth and seventh variants replace the AWD-LSTM encoder with transformers of varying sizes. “RAPPPID+TransfLG” has six layers and eight heads and feed-forward networks with 2048 dimensions, while “RAPPPID+TransfSM” has fewer parameters (two layers, two heads, and 80 dimensions). Both transformer variants have a dropout rate of 20%. The number of heads and layers in RAPPPID+TransfLG were chosen to closely match those used in the BERT language model (Devlin *et al*., 2019) whose architecture has been previously used for predicting protein function (Elnaggar *et al*., 2021). The number of heads and layers for RAPPPID+TransfSM were chosen to be representative of reasonable lower bounds for those hyper-parameters. The results of these analyses (Table 2) reveal that both of these models have a worse performance compared to the original RAPPPID model. The superior performance of the small transformer compared to the large one suggests that the large number of parameters required by transformers are leading to severe overfitting in this task and are the reason behind the performance deterioration

### Transfer Learning on Protein-Ligand Data from X-Ray Crystallography Experiments

RAPPPID’s strong performance on C3 datasets demonstrate that it is capable of generalising to predictions between proteins absent from the training set. Additionally, the ability to generalise these models to make predictions on test datasets that are not representative of the training data (e.g., different types of experiments or protocols are used to generate them) is also often desirable, yet is often much more challenging. Transfer learning is one approach that can be used to overcome the challenges imposed by differences between training and test datasets. Transfer learning is the process by which a network is first trained on a large, high-quality dataset. Many of these weights are then “frozen” such that they are no longer backpropagated through, while the remaining “unfrozen” weights are trained on a dataset that is often smaller and/or belong to a different input distribution (Yosinski *et al*., 2014). Here, we use this approach to tackle an interaction prediction task on a dataset that differs fundamentally from STRING: protein-ligand interactions and sequences as determined from X-ray crystallography data.

STRING and similar PPI datasets curate interactions from a wide range of modalities and provide sequences from reference proteomes. We’ve constructed a protein-ligand interaction task by leveraging BioLiP, a semi-manually curated list of X-ray crystallography experiments recorded in the Protein Data Bank (PDB) which reflect interactions between proteins and ligands (Yang *et al*., 2012; Berman, 2000). Using these datasets, we were able to extract sequences from the PDB records and, after filtering out interactions with low-quality sequences, construct a novel protein-ligand dataset. This dataset is available for download at https://github.com/jszym/rapppid. The sequences extracted from PDB are measured by fundamentally different modalities from those present in STRING/UniprotKB, and are often truncated and incomplete as a result. Furthermore, the nature of the interactions recorded by BioLiP are fundamentally different from STRING. Whereas STRING catalogues interactions between large classes of proteins, the nature of X-ray crystallography biases captured interactions to those with slower molecular dynamics and which do not primarily exist in aliphatic environments (Carpenter *et al*., 2008). These fundamental differences between BioLiP and STRING make them ideal datasets to illustrate the ability to use transfer learning with RAPPPID.

We first pre-trained RAPPPID on data from STRING, and then fine-tuned it on the BioLiP dataset after the LSTM encoder weights were frozen, leaving only the fully-connected classifier to be trained. To ensure no data leakage between the STRING and BioLiP datasets exist, protein sequences in STRING determined to have more than 90% identity with those in BioLiP were moved from the pre-training dataset to the fine-tuning dataset. C3-type training, validation, and testing splits were constructed for both the pretraining and fine-tuning datasets for evaluation purposes. Using this approach, RAPPPID pre-trained on the STRING dataset and fine-tuned on a portion of the BioLip dataset achieved an AUROC of 0.909. To test our hypothesis that transfer learning is indeed necessary to achieve good performance on this dataset, we also directly applied the pre-trained RAPPPID model (without any fine-tuning on BioLip) to the BioLip test split. As expected, this resulted in almost random results (AUROC = 0.548). These results highlight the difficulties of generalising a model to fundamentally different datasets and emphasise the utility of transfer learning to achieve good performance using RAPPPID in such scenarios.

### RAPPPID predicts interaction of HER2 with Trastuzumab and Pertuzumab

Peptides and proteins have emerged as an important class of therapeutics, enabling researchers to target previously “undruggable” targets (Tsomaia, 2015). Hundreds of peptide and protein therapeutics have been approved by the U.S. Food and Drug Administration (Usmani *et al*., 2017) with applications in treating illnesses ranging from cancer to heart disease (Sikder *et al*., 2019; Chen *et al*., 2012). Here, we sought to illustrate how one might use RAPPPID to validate hypothesised interactions between target proteins and candidate therapeutic proteins and peptides through two examples: Trastuzumab and Pertuzumab.

Trastuzumab and Pertuzumab are two recombinant humanised monoclonal antibodies which are used in combination in the treatment of metastatic breast cancers which belong to the HER2-positive subtype (Boekhout *et al*., 2011; Malenfant *et al*., 2014). Both Trastuzumab and Pertuzumab target distinct domains of the human epidermal growth factor receptor 2 (HER2). We applied RAPPPID, trained on our *H. sapiens* STRING C2 dataset (which is more appropriate for this application), to the sequences of the Trastuzumab and Pertuzumab antibody chains. RAPPPID predicted that HER2 interacts with Trastuzumab and Pertuzumab with 86.20% and 95.11% probability, respectively. This suggests that RAPPPID may be utilized as part of an interaction-based low-cost filtering or early validation step in the development of therapeutic proteins and peptides.

## Discussion and Conclusion

This study introduced RAPPPID, a deep learning method that addresses the challenges of creating generalisable PPI prediction models posed by inherent characteristics of PPI datasets. By adopting a modified AWD-LSTM training routine, RAPPPID was able to surpass state-of-the-art models under testing conditions that carefully controlled for information leakage and other sources of prediction accuracy inflation. Further experiments were conducted to confirm the results were independent of the specific proteins present in the training and testing splits. RAPPPIDs ability to PPIs in the STRING database was shown to increase with strong biological evidence for the interaction. This relationship between PPI evidence and RAPPPID predictive ability illustrates that RAPPPID accurately reflects our confidence in interactions, and testing performance is not disproportionately inflated by spurious, low-confidence interactions. Moreover, assessment of the sequence similarity between testing and training proteins revealed that the superior performance of RAPPPID is not due to the presence of highly similar protein pairs in testing and training, and the accuracy of RAPPPID was largely stable with a small improvement when highly similar testing proteins were excluded.

Developing appropriate and meaningful benchmark datasets for PPI prediction remains a challenging problem for a number of reasons. Firstly, deep learning tasks rely on large, high-quality datasets to obtain meaningful generalised models. Such datasets are few and far between. Projects like HIPPIE and iRefWeb join STRING in being among the best examples of PPI datasets which integrate multiple sources to assure both quality and quantity of PPI edges (Alanis-Lobato *et al*., 2017; Turner *et al*., 2010). Despite this, STRING is the only dataset among these three which has an appropriate number of high-confidence edges for the purpose of learning deep learning model. Specifically, there are 98.5% fewer edges in HIPPIE than in STRING between human proteins at a 95% confidence threshold. Even when confidence thresholds are lowered to 85%, HIPPIE holds 87.9% fewer human edges than STRING. Similarly STRING at a 95% confidence threshold has 75% more human edges when considering iRefWeb edges with 3 supporting references or more.

Secondly, this reliance on large, representative datasets is further exacerbated by the unique overfitting challenges posed by the characteristics of PPI data, as explored in this work and in (Park and Marcotte, 2012). While RAPPPID mitigates the overfitting tendencies of PPI data through regularisation, it also relies on large, representative training data to do so. In the construction of our C3 datasets, it is necessary to discard edges which are not between proteins of the same validation split. As a result, we observe a further decrease of up to 33.9% in the number of the already-precious-few edges afforded to us by STRING.

As we’ve shown in this work, the construction of benchmark datasets is critical for effectively comparing and evaluating PPI prediction tasks. It is thanks to large, quality PPI dataset projects like STRING that we might construct meaningful and appropriate datasets against which to evaluate PPI prediction methods like RAPPPID. Additional efforts and methodologies to collect and integrate PPI edges of the scale of STRING at high confidence levels are greatly desired, as they can help identify and mitigate biases that inevitably arise when constructing datasets.

The task of PPI prediction is related to the problem of protein docking inference, whereby computational models predict the atomic interactions between two proteins. While many methods have historically been based on the fast Fourier transform for energy evaluation (Desta *et al*., 2020), new and effective deep learning methods have also become available. One such method is AlphaFold-multimer (Evans *et al*., 2021), which builds upon a Transformer model for the prediction of protein structure (Jumper *et al*., 2021) to infer the interface of homo and hetero protein dimers at an atomic level. Integrating docking predictions can be an interesting future direction to help improve RAPPPID’s PPI prediction generalisation.

RAPPPID’s ability to predict interactions warrants further study into relevant tasks that might benefit from a similar approach. The RAPPPID architecture might be modified for the tasks of binding site and protein function prediction. These tasks are related to PPI prediction and as a result are exposed to similar challenges to which RAPPPID is well suited. However, in all these cases, it is crucial to consider strict rules for cross-validation and data splitting to ensure data leakage is avoided.

## Authors’ contributions

AE and JS conceived the study, designed the project and the algorithm, and wrote the manuscript. JS implemented the pipeline and performed the statistical analyses of the results. All authors read and approved the final manuscript.

