## Supplementary Material for "RAPPPID: Towards Generalisable Protein Interaction Prediction with AWD-LSTM Twin Networks"

Joseph Szymborski[1,2] and Amin Emad[1,2,3,*]

[1] Department of Electrical and Computer Engineering, McGill University, Montréal, QC, Canada

[2] Mila, Quebec AI Institute, Montréal, QC, Canada

[3] The Rosalind and Morris Goodman Cancer Institute, Montréal, QC, Canada

* Corresponding Author:

Amin Emad

755 McConnell Engineering Building

3480 University Street

Montréal, QC, Canada, H3A 0E9

**Supplemental Tables**

**Supplementary Table S1:** Ranges of hyperparameters used for training of RAPPPID. This table shows the range of hyperparameters and their increments considered when training RAPPPID. The final values of hyperparameters were chosen based on the performance on a randomly selected subset of the training set (i.e., a validation set).

| Hyperparameter | Number of LSTM Layers | Embedding Dropout Rate | LSTM Dropconnect Rate | Classifier Dropout Rate | Learning Rate |
|---|---|---|---|---|---|
| Range | 2-3 | 0.1-0.4 | 0.1-0.4 | 0.1-0.4 | $10^{-3}$ - $10^{-2}$ |
| Increment | 1 | 0.1 | 0.1 | 0.1 | 0.009 |

**Supplementary Table S2:** The final values of hyperparameters selected for the analysis reported in Table 1 of the manuscript for different datasets. The hyperparameters are selected based on the performance of a validation set (separate from the testing set).

| Experiment | Number of LSTM Layers | Embedding Dropout Rate | LSTM Dropconnect Rate | Classifier Dropout Rate | Learning Rate |
|---|---|---|---|---|---|
| C1 | 3 | 0.1 | 0.1 | 0.1 | $10^{-2}$ |
| C2 | 2 | 0.3 | 0.3 | 0.2 | $10^{-2}$ |
| C3 | 2 | 0.3 | 0.3 | 0.2 | $10^{-2}$ |

**Supplementary Table S3:** Statistics for the STRING C1, C2, and C3 datasets.

| | C1 | C2 | C3 |
|---|---|---|---|
| Number of Proteins | 9340 | 9340 | 9279 |
| Number of Edges | 263130 | 263130 | 173914 |
| Training Split Size (%) | 79.9% | 64.1% | 97.0% |
| Validation Split Size (%) | 10.0% | 19.0% | 1.6% |
| Testing Split Size (%) | 10.1% | 16.9% | 1.4% |
| Train/Validation Protein Overlap (%) | 100% | 83.2% | 0% |
| Train/Test Protein Overlap (%) | 100% | 84.3% | 0% |
| Train Edges w/ Test Proteins | 9137 | 3758 | 0 |
| Train Edges w/ Validation Proteins | 9170 | 3663 | 0 |

### Supplementary Figures

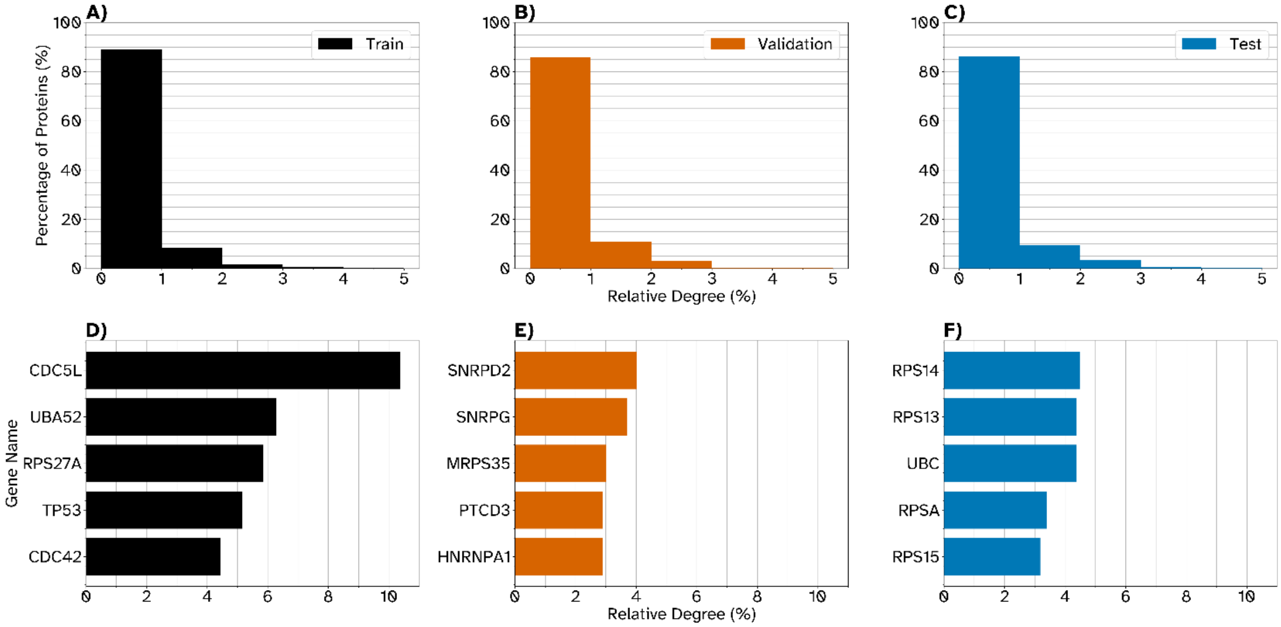


**Figure S1: Relative Degree of Proteins in the *H. sapiens* STRING C3 dataset.** The distribution of the relative degree of proteins between the three splits of the *H. sapiens* STRING C3 datasets reveal that the vast majority of proteins have a relative degree of less than 1% (A-C). The top five proteins with the highest relative degrees are plotted in bar graphs D through F. CDC5L has a relative degree of 10.37%, the highest of the entire dataset. Many of the proteins of high relative degree belong to the families of ribosomal or ribonucleoproteins.

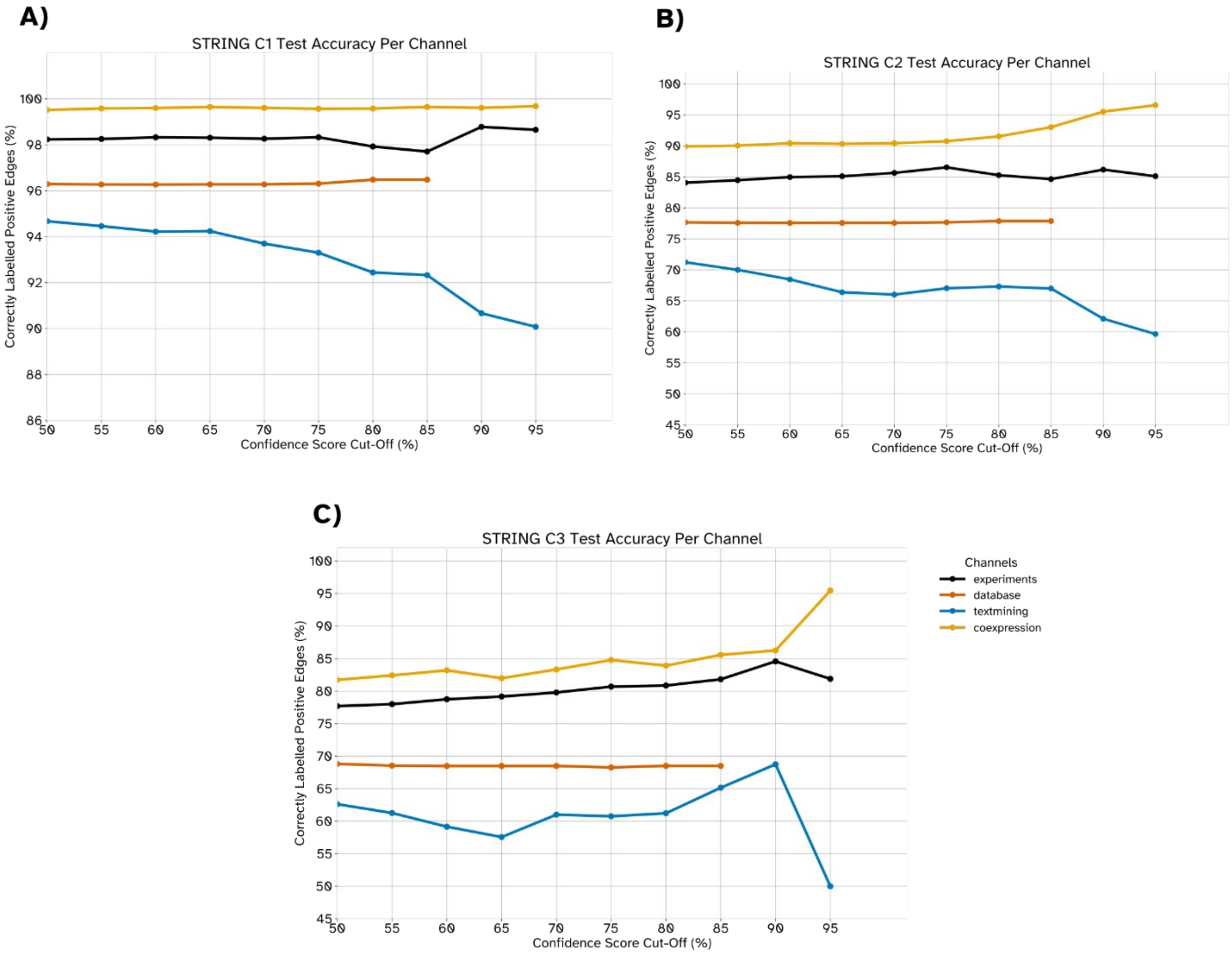


**Figure S2: Accuracy of positive edges across edge confidence stratified by STRING channels.** The percentage of correctly labelled positive edges are plotted for each major STRING channel in the C1 (A), C2 (B), and C3 (C) datasets. The x-axis denotes the channel edge confidence cut-off score for each curve's respective channel.

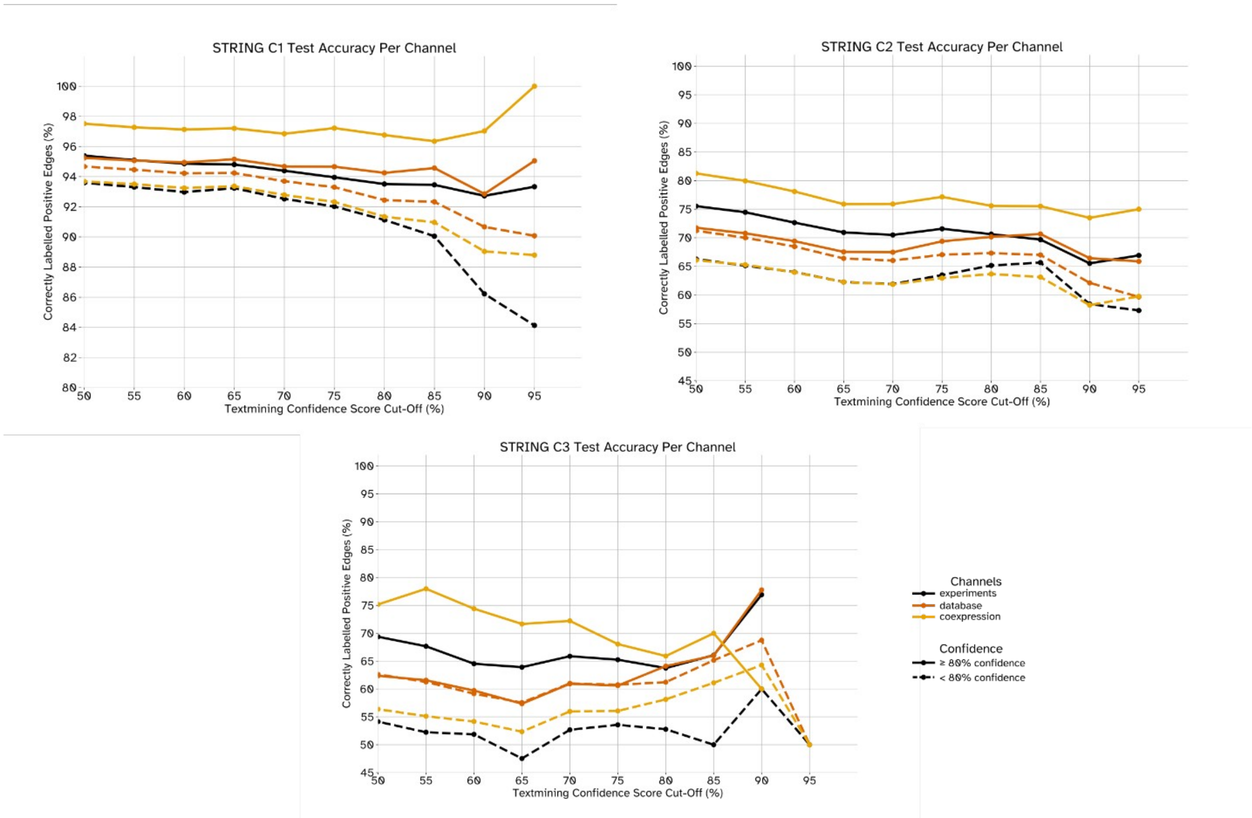


**Figure S3: Accuracy of positive edges across edge confidence stratified by STRING channels.** The percentage of correctly labelled positive edges are plotted for each major STRING channel, excluding text-mining. Edges are excluded according to the text-mining confidence threshold (x-axis). The solid curves include edges which have a channel confidence $\geq 80\,\%$ for the channel indicated by the curve's colour. Dashed curves conversely include edges whose channel confidence is ¿80 % for the channel indicated by the curve's colour. Data shown reflects the C1 model/dataset in panel (A), the C2 model/dataset in panel (B), and the C3 model/dataset in panel (C).

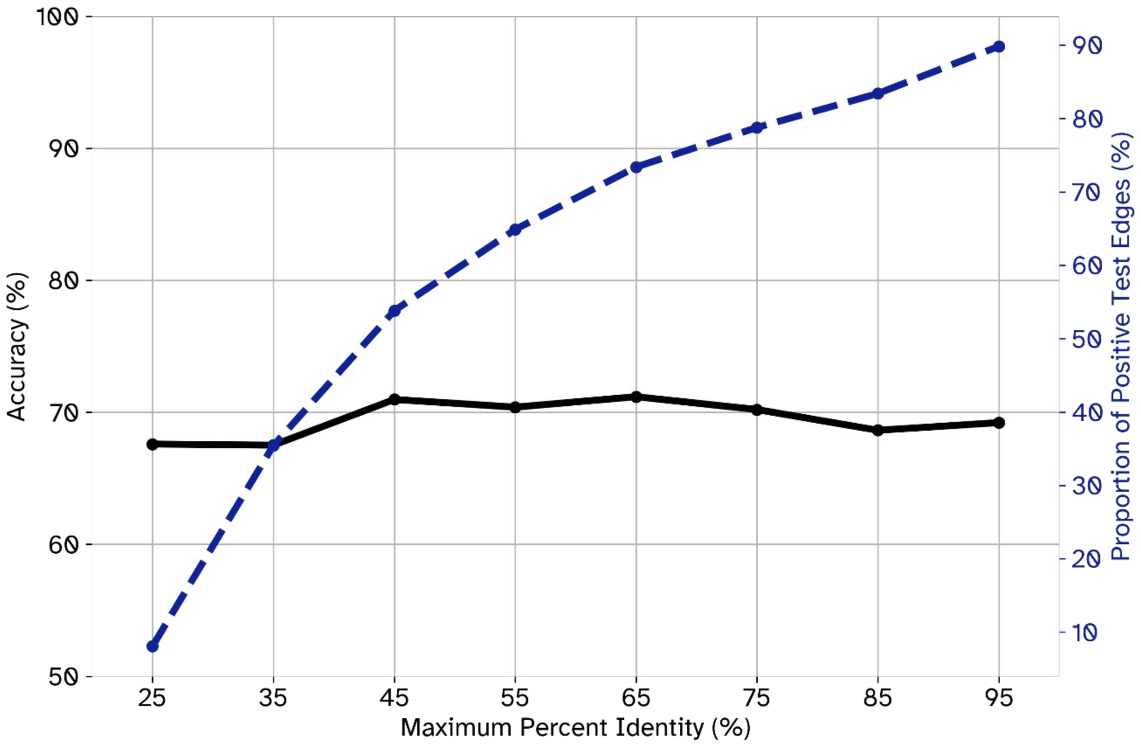


**Figure S4: Accuracy of positive edges as a function of similarity between testing and training proteins in C3.** The similarity between testing and training proteins was measured using their percent identity as computed by NCBI's PSI-BLAST software. The highest percent identity between any training protein and a testing protein in a testing edge was considered to be that testing edge's "maximum percent identity". The percentage of accurately labelled positive edges (black curve, left y-axis) is reported for edges with maximum percent identities lower than the threshold reported on the x-axis. The proportion of testing edges for each threshold values is reported by the dashed blue curve and the right y-axis.
